# Core architecture of a bacterial type II secretion system

**DOI:** 10.1101/397794

**Authors:** Anastasia A Chernyatina, Harry H Low

**Author notes:** Corresponding author: Harry H. Low, Department of Life Sciences, Imperial College, London, SW7 2AZ, UK.

## Abstract

Bacterial type II secretion systems (T2SS) translocate virulence factors, toxins and enzymes across the cell outer membrane (OM). An assembled T2SS has not yet been isolated in vitro. Here we use a fusion of negative stain and cryo-electron microscopy (EM) to reveal the core architecture of an assembled T2SS from the pathogen *Klebsiella pneumoniae*. We show that 7 proteins form a ∼2.5 MDa complex that spans the cell envelope. The outer membrane complex (OMC) includes the secretin PulD with all domains modelled and the pilotin PulS. The inner membrane assembly platform (AP) components PulC, PulE, PulL, PulM and PulN have a relative stoichiometric ratio of 2:1:1:1:1, respectively. The PulE ATPase, PulL and PulM combine to form a flexible hexameric hub. Symmetry mismatch between the OMC and AP is overcome by PulC linkers spanning the periplasm with PulC HR domains binding independently at the secretin base. Our results show the T2SS to have a highly dynamic modular architecture with implication for pseudo-pilus assembly and substrate loading.

## Introduction

The bacterial T2SS is found in human pathogens such as *Acinetobacter baumannii*, *Chlamydia trachomatis*, *Escherichia coli* and *Vibrio cholerae*^1^. It secretes a broad repertoire of substrates including digestive enzymes and infective agents like the cholera and heat-labile (LT) toxins^2^. Between 12-15 genes in a single operon usually encode the majority of T2SS components. Whilst the soluble domains for many of these proteins have been solved by X-ray crystallography^3,4^, their relative stoichiometry, mode of association with binding partners, and temporal coordination for assembling a functional secretion apparatus is still poorly understood.

The protein GspD forms a 15-fold rotationally symmetric pore termed the secretin that inserts into the OM and provides a conduit for substrate into the external environment. OM insertion is usually dependent on a lipidated pilotin^5^, which binds to the GspD C-terminal S-domain with 1:1 stoichiometry^6, 7^. The pilotin gene is often chromosomally discrete from the main T2SS operon. Multiple recent high-resolution cryo-EM structures report the partial secretin architecture^8-10^, and in complex with the pilotin^7^. However, the entire secretin has not yet been fully resolved due to disorder in the periplasmic N0 and N1 domains.

Within the inner membrane (IM), GspL and GspM are bitopic and monotopic membrane proteins, respectively, that together form homo- and hetero-dimers^11^. Combined with the polytopic membrane protein GspF^12^ and the ATPase GspE, these proteins form an assembly platform (AP) for the pseudo-pilus^13^. The pseudo-pilus constitutes a helical filament that extrudes fully-folded substrate through the secretin channel. The relative stoichiometry and overall ultrastructure of the AP is unknown. GspE is a cytoplasmic AAA+ ATPase that energises the T2SS and drives pilin assembly and pseudo-pilus formation. The active state is considered to be a hexameric ring as ATP turnover is significantly upregulated in an artificially oligomerized GspE-Hcp1 fusion^14^. The homologous ATPases PilB and PilT in the closely related type IV pilus (T4P) system also function as hexamers^15, 16^. Ultimately, the functional oligomeric state and stoichiometry of GspE within the T2SS apparatus has not yet been determined. The GspE N-terminal N1E domain connects to the N2E domain with an extended linker. A shorter but known flexible linker connects the N2E domain to the C-terminal CTE ATPase domain^17^. Such inherent flexibility within GspE is predicted to facilitate large-scale conformational changes. The N1E domain of GspE forms a 1:1 stoichiometric complex with the cytoplasmic domain of GspL^17, 18^. These two proteins contact GspF^13,19^, which is predicted to reside centrally within the AP. Concerted interplay between the GspE, GspL and GspF complex are thought crucial for coupling GspE conformational changes to the mechanical loading of pilin subunits within the pseudo-pilus assembly ^20, 21^. As GspL is a bitopic membrane protein, the direct contact between GspE and GspL also represents a mechanism for enabling cross-talk across the inner-membrane to other periplasmic components such as GspM. The coupling of the AP and OMC across the cell envelope is mediated by GspC, where the GspC N-terminus associates with GspL and GspM within the inner membrane^11^. The C-terminus of GspC must then span the periplasm as the GspC HR domain binds the GspD N0 domain^22^. Despite the known interaction between GspC HR domain and GspD N0 domain, the precise arrangement of GspC HR domain at the base of the secretin remains unclear^22^. Similarly, understanding how GspC HR domains bind to the secretin is important for determining how the likely symmetry mismatch between AP and OMC components is overcome.

Here we isolate an assembled T2SS so that both OM and IM components are captured together. Using a fusion of cryo and negative stain EM, combined with stoichiometry measurements, we provide a reconstruction of the entire OMC and a model for the cytoplasmic components of the AP. Combined they reveal a glimpse at the core ultrastructure of this cell envelope spanning nanomachine.

## Results

### Purification, EM and stoichiometry of Pul_CDELMNS_

The T2SS from the human pathogen *K. pneumoniae* HS11286 strain comprises 13 genes in a single unidirectional operon termed PulC through to PulO (Figure 1A). Note that Pul and Gsp nomenclature relate to equivalent proteins in homologous T2SS systems. The pilotin PulS is located in a separate position within the chromosome. These 14 genes were cloned and over-expressed in *Escherichia coli*. Using affinity chromatography tags positioned on the cytoplasmic ATPase PulE and the periplasmic pilotin PulS, a complex containing 7 components was purified by two successive pulldowns. Glutaraldehyde stabilization was included after the initial pulldown. The complex comprised PulC, PulD, PulE, PulL, PulM, PulN and PulS, and is here termed Pul_CDELMNS_ (Figure 1B). SDS-PAGE band identification was confirmed by LC-MS/MS. It is unclear why PulF and pseudo-pilus components were not co-purified with Pul_CDELMNS_. Their inclusion may require intact membrane for stabilization. Co-expression of PulA substrate or *pulG* knockout did not promote their inclusion. Trace quantities of GspJ and GspK were identified on the gel by LC-MS/MS suggesting the pseudo-pilins were at least partially expressed. Visualisation of Pul_CDELMNS_ by negative stain EM yielded particles ∼40 nm long and 17-22 nm wide (Figure 1C). The OMC PulD secretin was readily identifiable within 2D class averages. Hanging beneath the OMC and separated by a 5-10 nm gap the inner membrane AP was observed. Highly flexible linkers connect the OMC and AP so that these two assemblies effectively constitute independent particles tethered together. The Pul_CDELMNS_ complex was vitrified on thin carbon film and imaged by cryo-EM (Figure 1C). 2D class averages of Pul_CDELMNS_ yielded well resolved side views of the OMC. All domains of PulD were identifiable with additional densities observed at the base of the secretin where the PulC HR domain was expected to bind to the N0 domain^22^, and where PulS decorates the exterior of the secretin core^7^ (Figure 1C). The AP was not resolved here due to high flexibility and averaging effects. In the absence of glutaraldehyde, the OMC sometimes separated from the AP and yielded top views, which confirmed PulD and PulS in 1:1 stoichiometric ratio^7^ with C15 symmetry (Figure 1C). The relative stoichiometry of the inner membrane AP components were determined by SDS-PAGE densitometry and quantification of fluorescent emission using both Coomassie R250 and Sypro Ruby dyes^23, 24^ (Figure 1B). Pul_C:E:L:M:N_ relative mean ratios standardized around PulE were observed as 2.13:1.00:0.97:0.99:0.95, which is consistent with an overall relative ratio of 2:1:1:1:1 for these components. The relative ratios of PulE and PulL were tightly correlated (sigma = 0.07), which is important as it reflects their known 1:1 interaction ^17, 18^ and acts as an internal control for the gel densitometry. PulC stoichiometry was at least twice that of the other AP components with a relatively broad variance (sigma = 0.19). The 15-fold copy numbers of PulD or PulS were not used as a reference to determine an overall copy number for the AP components within the Pul_CDELMNS_ complex. PulD remained partially multimerized despite phenol treatment so that it failed to consistently enter and migrate through the gel fully. PulS quantity was significantly enriched as a consequence of the PulD secretin decoupling from the AP in the absence of glutaraldehyde stabilization during purification.

**Figure 1.**
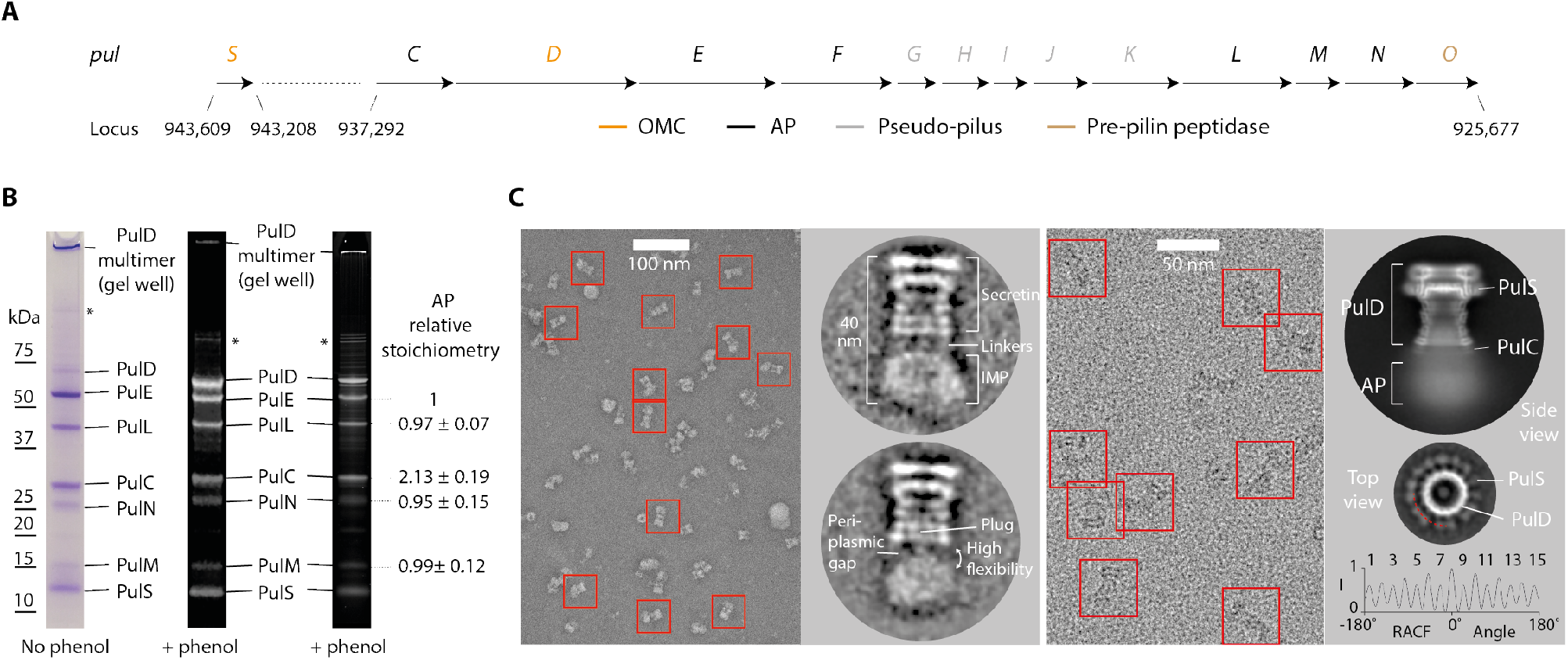
Purification, stoichiometry and EM analysis of the Pul_CDELMNS_ complex. **(A)** Schematic showing the chromosomal location and gene arrangement for the *Klebsiella pneumoniae* T2SS. OMC-outer membrane complex, AP - assembly platform. **(B)** Typical SDS-PAGE analysis of the Pul_CDELMNS_ complex. (Left) Coomassie R250 stained gel without phenol treatment. (Middle) Fluorescent emission Coomassie R250 stained gel imaged at 680 nm with phenol extraction. Residual PulD remains in the gel well. (Right) Fluorescent emission Sypro Ruby stained gel imaged at 302 nm with phenol extraction. Relative mean stoichiometry and standard deviation for all assembly platform (AP) components are indicated. Stoichiometry measurements were determined from six independent purifications. * indicates PulD multimer. **(C)** Left panel shows typical negative stain EM micrograph of the Pul_CDELMNS_ complex with selected individual particles highlighted with a red box. Zoomed images of 2D class averages are also shown. Right panel shows typical cryo-EM micrograph of the Pul_CDELMNS_ complex with associated side view 2D class average. Top view 2D class average shows PulS and PulD with C15 symmetry based on the RACF. Red dotted line indicates radial ring for RACF calculation.

### PulD secretin structure determination

Focused refinement of the OMC yielded a reconstruction with an overall resolution of 4.3 Å (Figure 2 and Figure S1). All domains of PulD were resolved with sufficient map quality (Figure S2A) to build a complete model of the monomer and the secretin, excluding the amino acids (aa) in loops 288-303, 462-470 and 632-637 (Figure 3 and 4A, and Figure S3). The PulD fold is similar to partial *Escherichia coli* K12 and H10407 GspD models (RMSD Cα = 3.2 Å and 3.3 Å) where the secretin core and N3 domain have been described, along with homology modelled N2 and N1 domains^7, 10^ (Figure 4B). The entire secretin is 20 nm long with an external diameter of 15 nm at the base (Figure 2B and Figure 3). It includes an occluding central gate and N3 domain constriction sites within the secretin channel. It lacks a *Vibrio cholerae* cap gate^10^. The N1, N2 and N3 domains pack tightly (Figure 5) with a diagonal offset of 36°. N0 is positioned almost directly below the N1 domain and does not maintain the diagonal offset. The N0 fold is similar to that described in multiple crystal structures^22,25,26^ with a core of two helices flanked on each side by β-sheets. However, its position relative to the N1 domain within the secretin is significantly different to these crystal structures where crystal contacts appear to have dominated domain arrangement (Figure S2B). The N0 and N1 domains are connected by loop 7, which constitutes a substantial 26 aa linker. The N-terminus of loop 7 forms a wedge that packs between neighboring N1 domains. Its C-terminus partially envelops the proximal N1 domain whilst making additional secondary contacts with N0 domain helix 2 (Figure 3A and Figure 5D). The N0 domains form a tightly packed ring with alternating stacked β-sheets sandwiched between helices 2 and 4 (Figure 5D). Failure to stabilize the N0 domain and to promote formation of the loop 7 wedge likely accounts for the previously reported N0, N1 and N2 domain flexibility in other systems^7-10, 27^. Overall, a single PulD monomer has a radial twist around the secretin long axis of 130° (Figure 3C).

**Figure 2.**
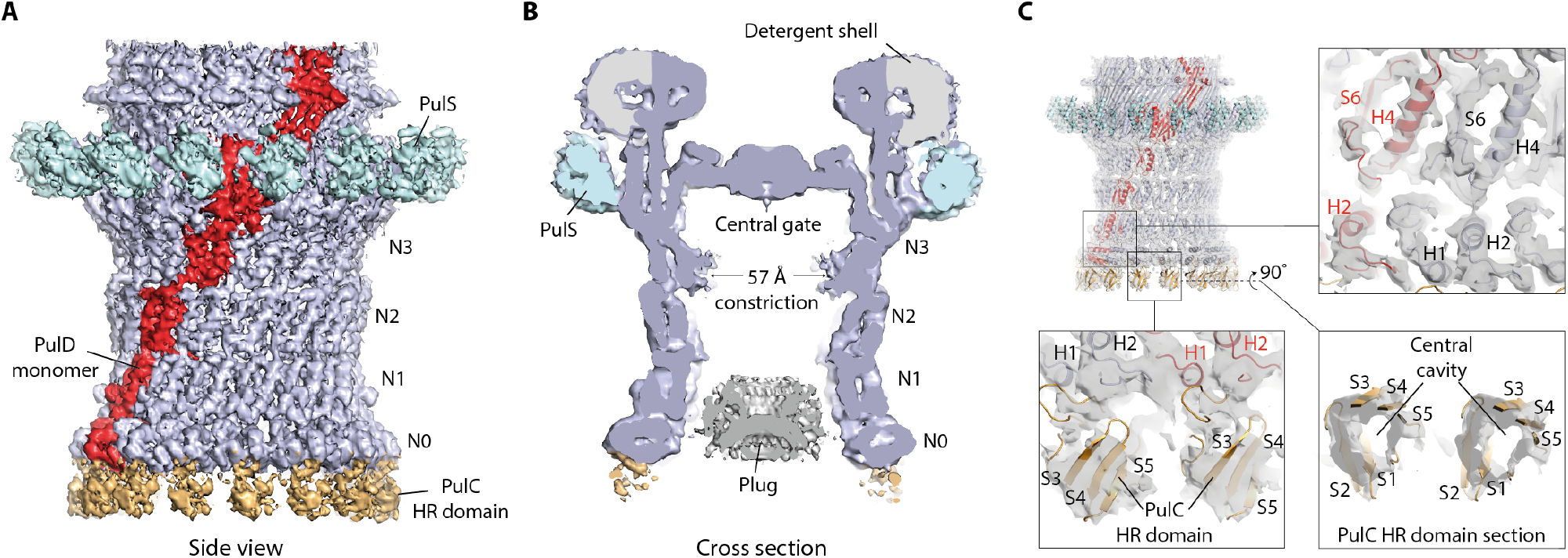
Cryo-EM density map of the outer membrane complex (OMC). **(A)** 4.3 Å resolution C15 symmetrized cryo-EM map of the OMC locally sharpened with LocScale^39^ and contoured at 5.5σ. **(B)** Cross section view of the OMC revealing internal features of the PulD secretin. A plug occludes the secretin lumen. The map was unsharpened and contoured at 3σ. **(C)** Fit of molecular models within the OMC map. *De novo* model for the PulD secretin including the N1 and N0 domains (top right). A PulC HR domain homology model was rigid body fitted. The map was sharpened with LocScale and contoured at 4σ (bottom left). In this localized region of the map at 7-8 Å resolution, the central cavity of the PulC HR domain was clearly resolved (bottom right).

**Figure 3.**
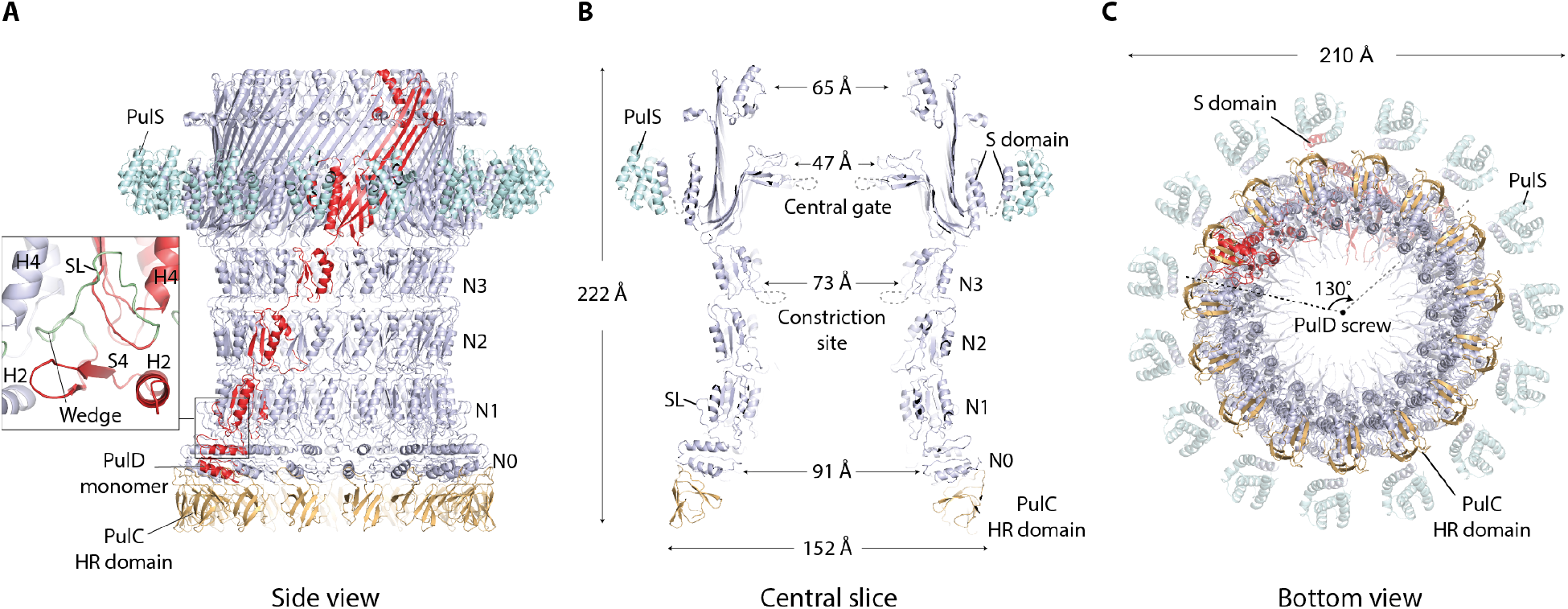
Structure of the outer membrane complex (OMC). **(A)** Cartoon representation of the OMC, which includes the C15 symmetrized secretin (light blue) with a single PulD monomer highlighted in red. PulS (cyan) and PulC HR domain (orange) homology models were fitted as rigid bodies. 15 PulC HR domains were modelled here although stoichiometry measurements indicate an occupancy of ∼0.5 in the Pul_CDELMNS_ complex. Zoom panel shows how loop 7 (green coil) constitutes a stabilizing loop (SL) that packs as a wedge between neighboring N1 domains. **(B)** Cross section view of the OMC structure with dimensions. **(C)** Bottom view of the OMC structure. Each PulD monomer has an azimuthal span twisting around the secretin long axis of 130°.

**Figure 4.**
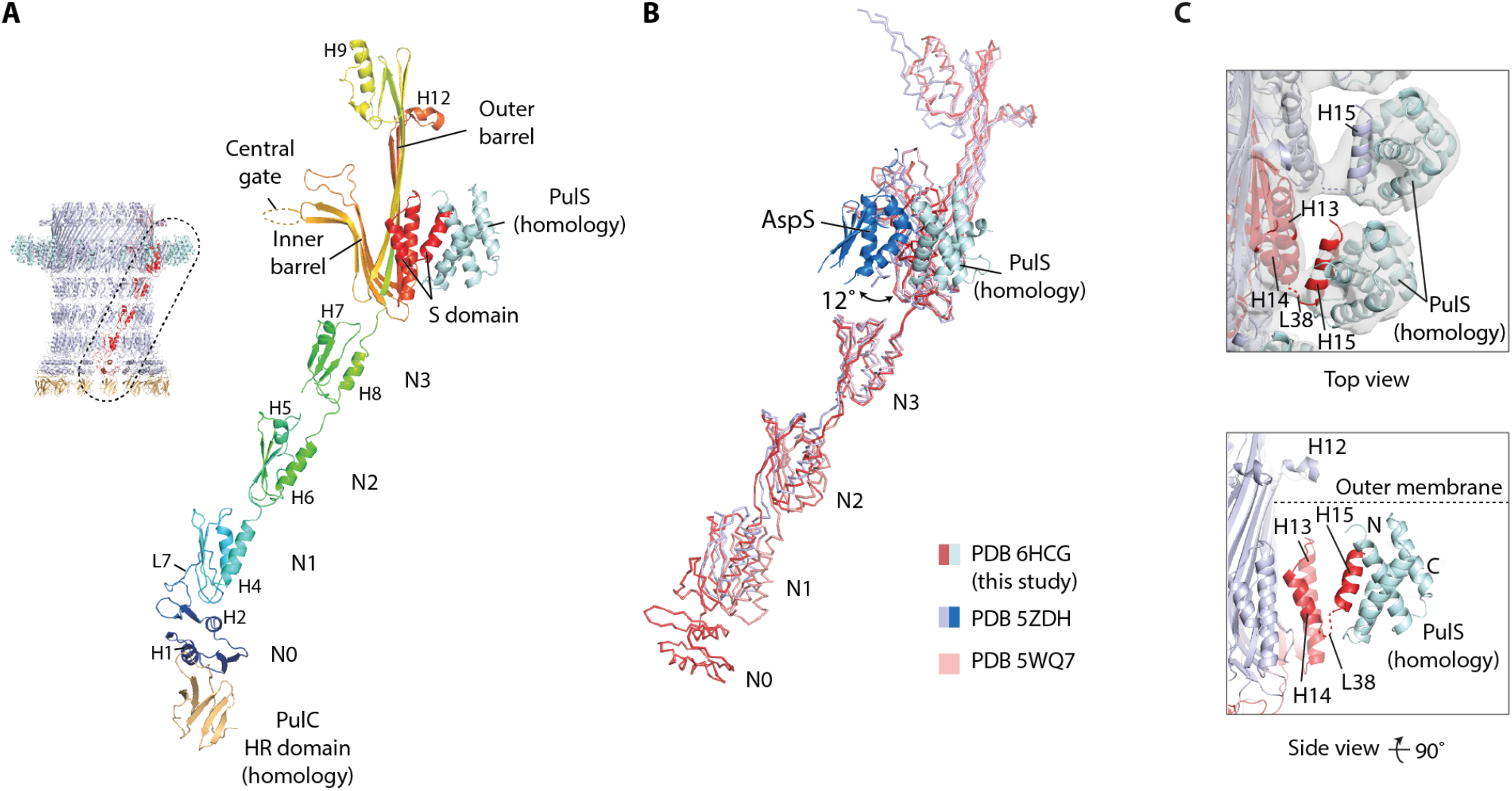
Structural comparison of PulD and PulS models. **(A)** A complex of PulD, PulS and PulC HR domain monomers extracted from the C15 OMC model (dotted region). For PulD, rainbow coloring highlights blue N-terminus through to red C-terminus. **(B)** Superposition of the PulD and PulS complex with the equivalent from enterotoxigenic *E. coli* PDB 5ZDH and *E. coli* K12 PDB 5WQ7. Compared to PulD, these PDBs differ by RMSD C*α* = 3.3 Å and 3.2 Å, respectively. The relative position of AspS in PDB 5ZDH differs to PulS by a 12° azimuthal rotation around the secretin long axis. **(C)** (Top) Fit of PulS homology model with the OMC map. The map was low pass filtered to 8 Å and contoured at 5σ before rigid body fitting. (Bottom) Cartoon schematic showing the PulS model relative to the PulD S-domain (red helices). PulD helix 15 locates to a groove within PulS^30^. Flexible loop 38 represents the only attachment between the helix 15/PulS complex and the rest of the PulD monomer.

**Figure 5.**
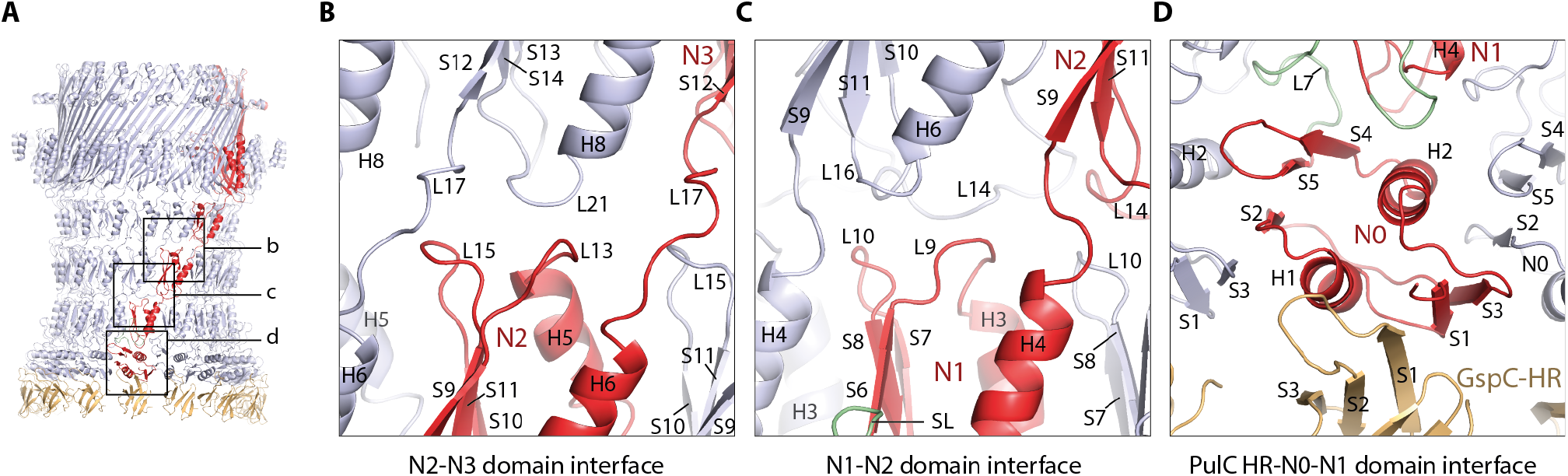
Structural analysis of the outer membrane complex (OMC) periplasmic domains. **(A)** Overview model of the OMC with PulS removed for clarity. Boxed regions are zoomed in **B-D**. **(B-D)** Cartoon representation of selected domain interfaces showing key structural details.

### PulC HR domain binds at the secretin base

Hanging beneath the N0 domains in the OMC map, additional globular densities at 7-8 Å resolution protrude from the secretin base (Figure 2 and Figure S1B). Focused refinements^28^ failed to markedly improve resolution. These densities were predicted to be the PulC HR domain given its known interaction with the PulD N0 domain in homologous systems^22, 29^. Using the GspC HR domain and GspD N0 domain crystal structure^22^ as a reference, a homology model of PulC HR domain was fitted as a rigid body (Figure S2C). The model closely follows the surface envelope of the map in this region, with a pair of triple β-sheets opposed around a central cavity (Figure 2C). Based on the quality of this fit, the PulC HR domain was assigned to each globular protrusion. Importantly, each PulC HR domain binds to only a single PulD N0 domain and has no contact with neighbouring PulC HR domains. Overall, the binding of the PulC HR domain to the PulD N0 domain appears to be important for the correct positioning of the N0 domain within the secretin and the subsequent stabilization of the N1 and N2 domains. Additional stabilization is derived from a plug that occludes the lumen of the secretin at the level of the N0-N1 domains (Figure 2B). The plug was observed in both the 2D class averages (Figure 1B) and the 3D reconstruction. The high-resolution plug ultrastructure was not resolved due to a likely symmetry mismatch with the C15 averaged OMC. Attempts to resolve the plug structure through refinement using lower symmetries yielded reconstructions of low quality and resolution. Further studies will be required to resolve the source of the plug although the PulC PDZ domain is a speculative candidate given the position of the PulC HR domain at the base of the secretin.

### The PulS pilotin decorates the secretin core

Decorating the outside of the secretin core proximal to the PulD S-domain in the map, globular densities were observed in a position consistent with the pilotin AspS relative to GspD in ETEC^7^ (Figure 2). For these densities, map resolution was limited to ∼7 Å (Figure S1) and focused refinements^28^ did not markedly improve resolution. A homology model of the PulS pilotin in complex with the PulD S-domain helix 15 based on the homologous structure in *Dickeya dadantii* (PDB 4K0U^30^) was fitted as a rigid body (Figure 4C). Compared to AspS^7,31^, the position of PulS differs by a 12° radial rotation around the secretin long axis (Figure 4B). Loop 38 between S-domain helix 14 and helix 15 bound to PulS constitutes the lone contact point between the secretin core and PulS (Figure 4C). No additional contacts were observed in contrast to AspS-GspD where the secretin core helix α11 forms extensive secondary contact with the pilotin^7^. The lack of equivalent secondary contacts between PulS and PulD likely accounts for the apparent flexibility between these proteins and may be a distinguishing feature between the structurally discrete *Klebsiella*-type and *Vibrio*-type pilotins.

### PulC links the OMC to the inner membrane AP

Whilst the PulC HR domain binds to the base of the secretin, its N-terminus is located within the inner membrane AP^11^ so that PulC is predicted to span the periplasm and link the AP and OMC. To verify the presence and positioning of the PulC N-terminus within the AP, a hexahistidine tag was inserted after aa 61 where PulC was predicted to exit the inner membrane and enter the periplasm. Ni-NTA gold labelling showed beads localize exclusively to the AP and not the OMC (Figure 6A and Figure S4). Given the PulC HR domains bind to the base of the secretin, PulC therefore spans the periplasmic gap between the OMC and AP (Figure 1C).

**Figure 6.**
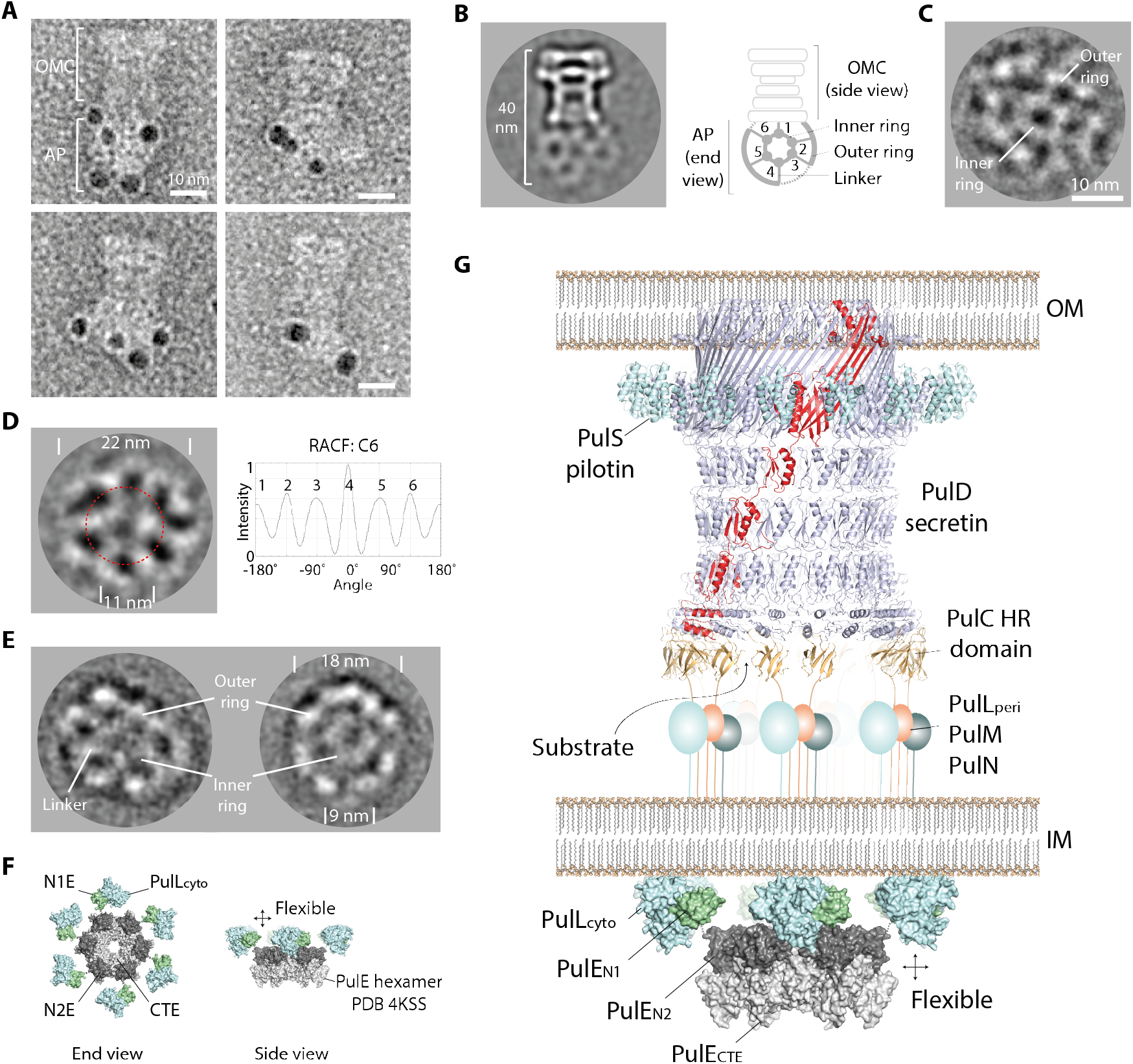
2D EM of the assembly platform (AP) and a T2SS core model. **(A)** Ni-NTA gold bead labelling of the PulC N-terminus. Beads locate exclusively to the AP. **(B)** Cryo-EM 2D class average of Pul_CDELMNS_. The AP showed end views as a preferred orientation with C6 symmetry. (**C)** Cryo-EM 2D class average of Pul_CDELMNS_ with the alignment and classification focused on the AP. **(D)** Cryo-EM 2D class average of Pul_ELM_ showing C6 symmetry. RACF = rotation auto-correlation function calculated around dotted red radial ring. **(E)** Negative stain EM 2D class average of Pul_ELM_. **(F)** Schematic showing modelled arrangement of the Pul_EL_ cytoplasmic domains within the Pul_CDELMNS_ and Pul_ELM_ hexameric hub. The central PulE hexamer (inner ring, PDB 4KSS) is decorated by six copies of the Pul_E-N1E/Lcyto_ complex (outer ring, PDB 2BH1). Connection between inner and outer ring is mediated by the 44 aa flexible N1E-N2E inter-domain linker. **(G)** Stoichiometric model of an assembled T2SS core apparatus. AP components comprise a hexameric hub. PulC connects the AP to the OMC by binding to the secretin base. Substrate may be loaded through the PulC cage in positions where GspC is absent.

### PulE, PulL and PulM form a flexible hexameric hub

Cryo-EM 2D class averages of the Pul_CDELMNS_ complex revealed the ultrastructure of the AP positioned beneath the OMC. A 20-22 nm outer ring is coupled to a 10-12 nm inner ring by six radial linkers (Figure 6B). Focused alignments of the AP where the OMC was masked out show the outer ring to be comprised of weakly associating non-contiguous globular densities (Figure 6C and Figure S5A). This concentric ring structure is highly flexible and represents the preferred single orientation of the AP so that 3D structure determination was impeded. The addition of non-hydrolysable ATP analogues made no obvious change to the Pul_CDELMNS_ complex under conditions tested. In order to dissect the observed AP ultrastructure, a sub-assembly constituting PulE, PulL, PulM, and PulN (Pul_ELMN_) was purified by 2-step affinity chromatography. GraFix^32^ stabilized the complex and reduced particle heterogeneity. PulN bound weakly to the complex and only trace quantities were observed by SDS-PAGE within the Pul_ELMN_ complex after GraFix (Figure S5B). The resultant Pul_ELM_ complex was analyzed by cryo and negative stain EM on continuous carbon film (Figure 6D-E and Figure S5B-C). Under vitreous conditions, Pul_ELM_ yielded a preferred orientation concentric ring structure that was similar to the AP within the Pul_CDELMNS_ complex, with equivalent dimensions and overall C6 symmetry. By negative stain, the same ultrastructure was observed although the sample was compacted so that the outer and inner rings have dimensions of 16-18 nm and 8-9 nm, respectively. Compaction was likely a consequence of sample flexibility and drying effects during the negative stain procedure. PulE alone, and PulE in complex with the cytoplasmic domain of PulL (aa 1-235) or full-length PulL were also purified from the membrane fraction. However, EM analysis yielded heterogeneous and flexible particles that did not further resolve AP ultrastructure under conditions tested.

## Discussion

### Model of the inner membrane AP

Negative stain and cryo-EM studies of the Pul_CDELMNS_ complex revealed the ultrastructure of the AP positioned beneath the OMC to be a highly flexible hexameric hub comprising two concentric rings. In vitreous ice, the inner ring had a diameter of ∼11 nm and overall C6 symmetry. The outer ring constituted non-contiguous globular densities with ∼22 nm diameter. Similar ultrastructure was observed with purified Pul_ELM_ complex meaning just these components alone are sufficient to generate the bulk of the hexameric hub structure. In both systems, the single preferred orientation captured was considered to represent an end view projection of the AP. In the T2SS, the *V. cholerae* PulE homologue GspE forms a quasi C6 ring with diameter 11-13 nm when combined with an Hcp1 fusion^14^. The related T4P system is energized by the ATPases PilB and PilT, which are known to operate as highly dynamic hexameric rings. PilB has dimensions of 13.5 nm and 9 nm when in an elliptical C2 conformation^15^ whilst the PilT ring has a diameter of 11.5 nm^33^. For PilB and GspE these dimensions are specific to the N2E and CTE ATPase domains and do not include their N-terminal N1E domains. In situ reconstruction of the T4P system in *Myxococcus xanthus* indicates the position of PilB and PilT to be located centrally at the base of the AP^34^. Collectively, this data is consistent with a model for the Pul_CDELMNS_ AP and Pul_ELM_ complex in which the observed inner ring constitutes a PulE hexamer comprising the N2E and CTE domains similar to that observed in *V. cholerae* GspE^14^ (Figure 6F).

The *V. cholerae* GspE N1E domain forms a direct 1:1 stoichiometric complex with the cytoplasmic domain of GspL^17, 18^. The equivalent domains in *K. pneumoniae* also form a 1:1 stoichiometric complex termed here Pul_E-N1E/Lcyto_ (Figure S5D). In the T4P system, the PulL_cyto_ homologue PilM forms a ring around the central PilB/PilT hexamer. This architecture supports a model for the Pul_CDELMNS_ AP and Pul_ELM_ complex where the bulk of the hexameric hub outer ring constitutes Pul_E-N1E/Lcyto_ complexes (Figure 6F). PulM, the periplasmic domain of PulL (termed PulL_peri_), and PulN and PulC when present, may also contribute to the observed outer ring densities if ordered. Certainly PulL_peri_, PulM and PulN do not form a periplasmic ring here as might be expected based on the T4P system^34^. The retention of membrane or inclusion of other T2SS components such as the pseudo-pilus may be required for the stabilization and visualization of any such ring.

Within the Pul_CDELMNS_ complex, the mean relative ratio of PulC:PulE: PulL:PulM:PulN was measured to be 2.13:1.00:0.97:0.99:0.95 indicating two copies of PulC for each of the AP components. The globular densities that comprise the outer ring of the hexameric hub do not form a contiguous well-ordered structure. This observation combined with the stoichiometry supports a modular architecture where independent or weakly associated sub-complexes containing two copies of PulC and a single copy of PulM, PulN, and Pul_E-N1E_ in complex with PulL decorate each subunit of the central PulE hexamer (Figure 6G). This model does not preclude the possible formation of a Pul_E-N1E/Lcyto_ ring positioned more centrally directly above the PulE N2E/CTE hexamer *in vivo*. Such an arrangement would be similar to that observed for the N1 domains of *Thermus thermophilus* PilF^35^ and may promote the close contact and encircling of the putative pilin spooling protein PulF^13, 19^.

### PulC mechanism for overcoming AP-OMC symmetry mismatch

PulC spans the periplasm to bind the OMC with the PulC N-terminus located within the inner membrane and the HR domain bound to the PulD N0 domain at the secretin base. Based on six PulE subunits within the AP, stoichiometry measurements support a model where the PulC copy number within the AP is 12. PulC therefore constitutes a cage that spans the periplasm (Figure 6G) and is reminiscent of the virB10 N-terminus which spans the periplasm in the type IV secretion system^36^. Substrate may have the potential to gain access to the secretin lumen via PulL and PulM^37^ in positions where PulC is absent. The OMC structure shows that each PulC HR domain binds a single PulD N0 domain independently with no lateral contacts between neighbouring HR domains. This is important as it provides a natural mechanism for overcoming symmetry mismatch between the 12 PulC subunits located within the hexameric AP and the 15-fold symmetric OMC. In this model on average three secretin N0 domains remain unoccupied without bound PulC HR domain. Whether full secretin N0 domain binding occupancy occurs with up to 15 bound PulC HR domains cannot be entirely excluded as each of the N0 domains has at least the potential to bind a PulC HR domain. T2SS assembly and secretion may ultimately occur in a dynamic equilibrium with a range of N0 domain binding occupancies.

### The secretin N0-N1 domains form a compact C15 ring

Multiple T2SS secretin structures have been solved including *Klebsiella*-types from *E. coli* K12^10^ and *Pseudomonas aeruginosa*^9^, and *Vibrio*-types from enterotoxigenic *E. coli* (ETEC)^7^, enteropathogenic *E. coli* (EPEC)^8^ and *V. cholerae*^10^. Despite high quality 3-3.5 Å resolutions achieved for the secretin core and N3 domains, all the structures have poorly resolved densities towards the PulD/GspD N-termini with homology modelling required for the N1 domains and entirely absent N0 domains. Docking of N0 and N1 domain co-crystal structures^22, 25-27^ into the secretin maps failed to resolve the possible position and orientation of the N0 domain due to significant steric clashes. Consequently, the PulC HR domain orientation relative to the N0 domain remained unclear and it was modelled decorating the inside of the secretin base^22^. To account for the high flexibility of the secretin N0 and N1 domains and the symmetry mismatch between the 6-fold symmetric AP and 15-fold OMC, a pseudo-6-fold arrangement comprising a hexamer of dimers had been proposed for the N0-N2 domains^9^. In such a model the N0-N2 domains of three secretin subunits would be displaced in a metastable arrangement. The OMC structure derived from the Pul_CDELMNS_ complex resolves now how the PulD N0 domains are organized within the secretin relative to the N1 domain ring. The N-terminus of loop 7 wedges and stabilizes neighbouring N1 domains whilst the N0 domains form an intimate and stable C15 ring. At least when PulC HR domain is bound, the N0 ring does not immediately support a model where the secretin incorporates multiple symmetries along its length. Instead the secretin forms a well-ordered barrel, which combined with the PulC HR domains bound at the base protrudes from the outer membrane ∼20 nm into the periplasm.

### Concluding remarks

In this study, the T2SS outer membrane components PulD and PulS are co-purified with the core components of the inner membrane AP including PulC, PulE, PulL, PulM and PulN. Overall, our results reveal a glimpse at the core architecture of an assembled T2SS (Figure 6G) and show it to be different to other known secretion systems (Figure S6). The T2SS AP does not constitute a stack of conjoined rings that seal directly to the secretin base forming an enclosed central channel as observed in the T3SS^38^. Instead, the T2SS has a highly dynamic and modular architecture where the OMC and inner membrane AP have weakly associating and limited inter-connection via a PulC cage. This observation is consistent with the requirement for substrate to gain access to the secretin lumen via the periplasm. Whilst the resolution of the hexameric hub observed in the Pul_CDELMNS_ and Pul_ELM_ complexes is limited it shows the T2SS to have a six-fold arrangement for assembled AP components. The observed AP architecture is consistent with that described in the T4P system^34^ providing evidence that the inner membrane ultrastructure is broadly conserved between these different but related secretion systems. Although the connection between inner and outer membrane complexes was expected to be transient and dynamic^3^, we show that it is possible to isolate an almost fully assembled T2SS *in vitro*. Our work suggests that the purification of an entire T2SS with the pseudo-pilus, PulF, and ultimately substrate incorporated is in reach. These additional components may be necessary to help stabilize the inner membrane AP so that ultimately full structural dissection and mechanistic understanding for the T2SS may be achieved.

## Acknowledgements

We thank eBIC for cryo-EM data collection support particularly Kyle Dent and Yuriy Chaban. Tillmann Pape and Paul Simpson for in-house EM support. Francesca Gubellini for gold labelling advice. Arjen Jakobi and Carsten Sachse for LocScale support. This work was funded by a Wellcome Trust Career Development Fellowship Enhancement Award (200074/Z/15/Z) to H.L.

## Author Contributions

A.C and H.L designed experiments. H.L initially cloned and purified the Pul_CDELMNS_ complex. A.C and H.L cloned and purified proteins, collected and processed data, and solved structure. A.C built the OMC structure. A.C and H.L. determined stoichiometry. A.C. undertook gold labelling. H.L wrote the paper with contributions from A.C.

## Data availability

3D cryo-EM density maps produced in this study have been deposited in the Electron Microscopy Data Bank with accession code EMD-0193. Atomic coordinates have been deposited in the Protein Data Bank (PDB) under accession code 6HCG.

## Supplementary Figures

**Figure S1.**
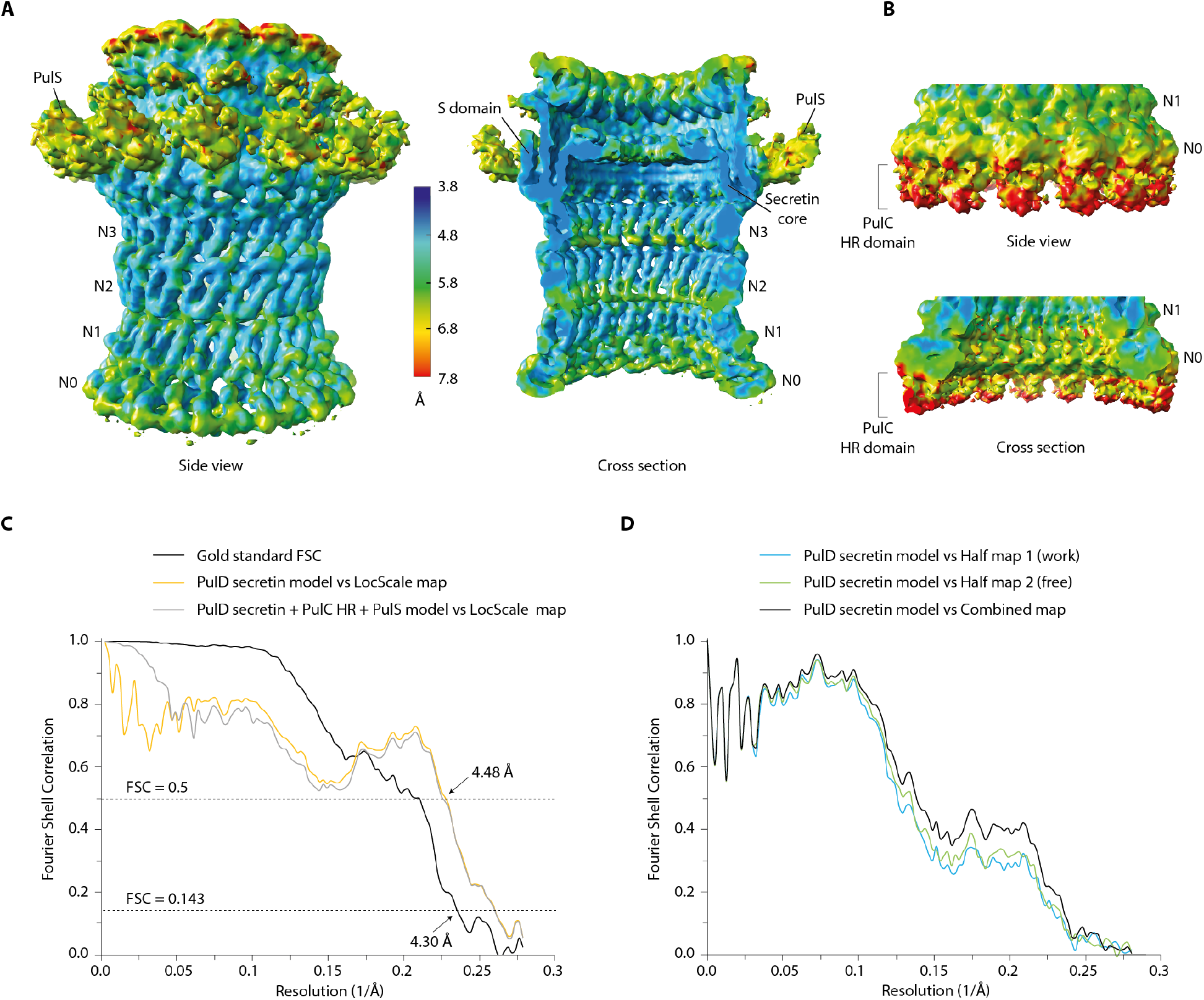
Outer membrane complex (OMC) local resolution map and FSC curves. **(A)** Unsharpened OMC map showing local resolution estimates calculated using ResMap^40^ and contoured at 4.5σ. **(B)** Equivalent to **A** but contoured at 2σ so as to view the PulC HR domain densities at the secretin base. **(C)** Gold standard FSC curve of the OMC (black). FSC curves between the PulD secretin model and the LocScale^39^ sharpened map (orange). FSC curves between the PulD secretin model with homology models of PulS and PulC HR domain fitted, and the LocScale sharpened map (grey). **(D)** Cross-validation for the PulD secretin model. PulD secretin model versus half map 1 (work-light blue), half map 2 (free-green), and combined (black). See Methods for further detail.

**Figure S2.**
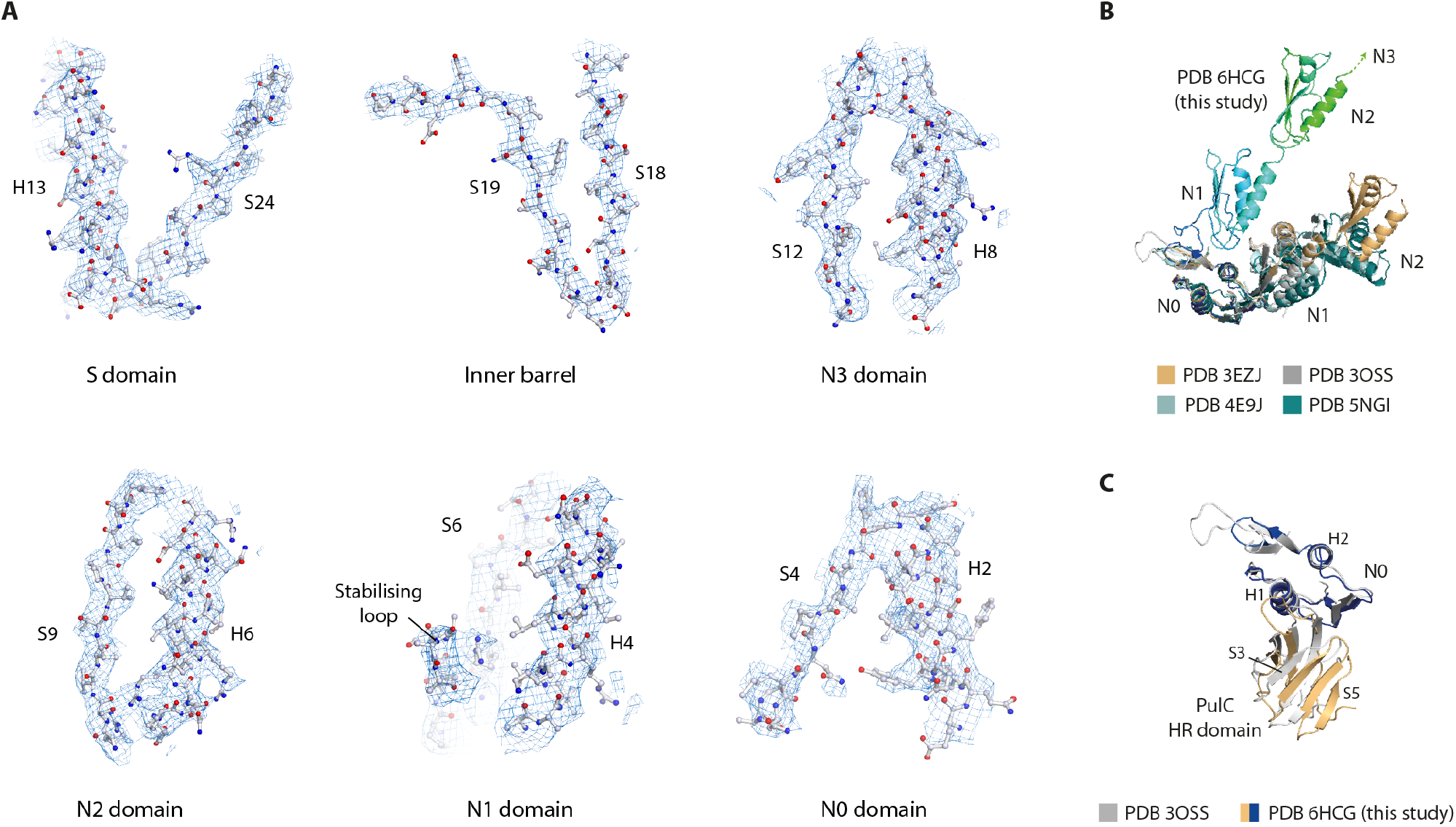
Map quality and PulS fit. **(A)** Selected regions of the OMC EM density map showing PulD secretin side chain detail and build. The map was sharpened with B factor = - 142 Å^2^ and contoured between 5-8σ. **(B)** Superposition of PulD N0, N1 and N2 domains with equivalent domains derived from various crystal structures including enterotoxigenic *E. coli* PDB 3EZJ and PDB 3OSS and *P. aeruginosa* PDB 4E9J and PDB 5NGI. **(C)** Superposition of PulD N0 domain in complex with PulC HR domain against the equivalent domains in the enterotoxigenic *E. coli* crystal structure PDB 3OSS. The complexes differ by RMSD C*α* = 1.4 Å.

**Figure S3.**
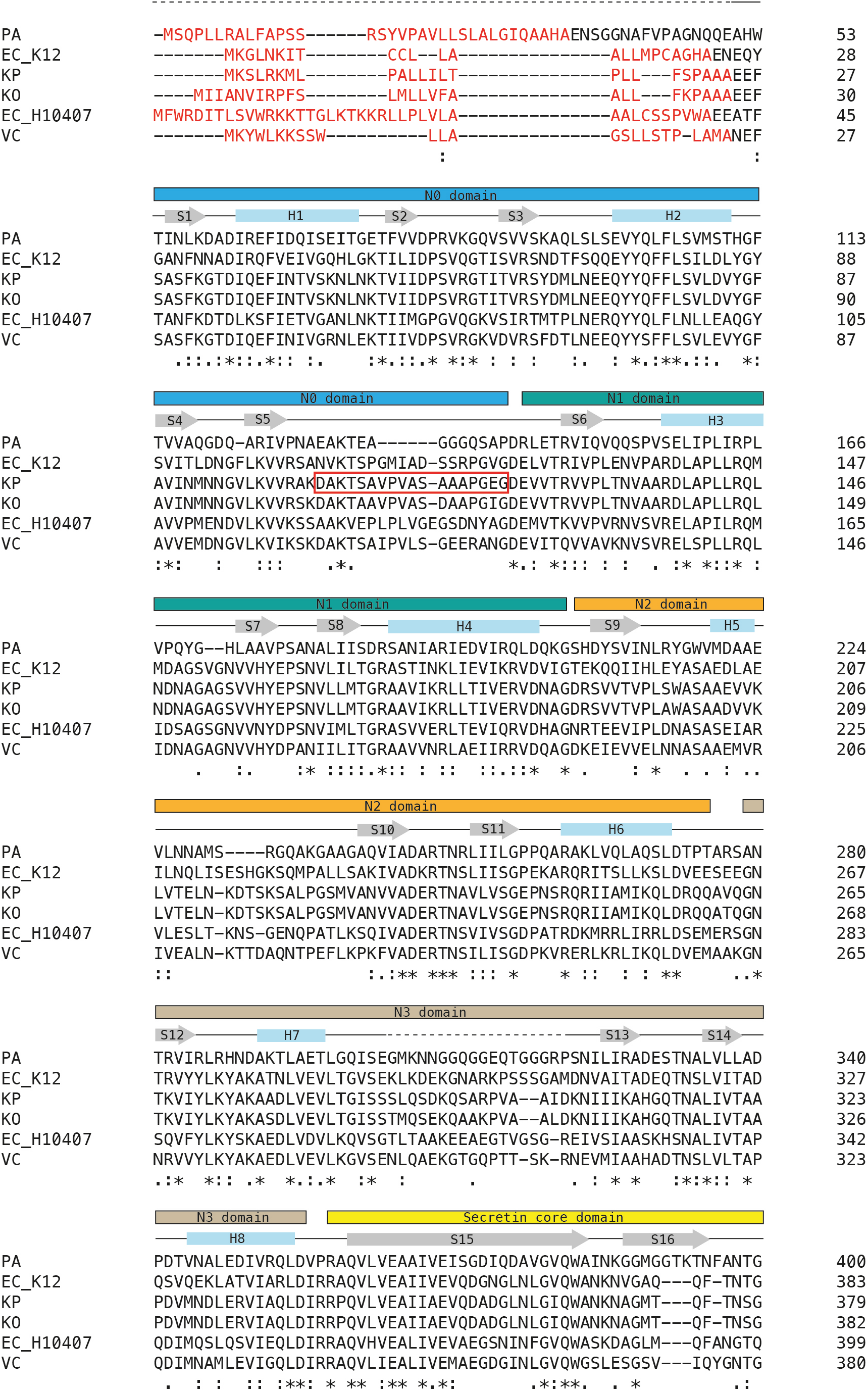
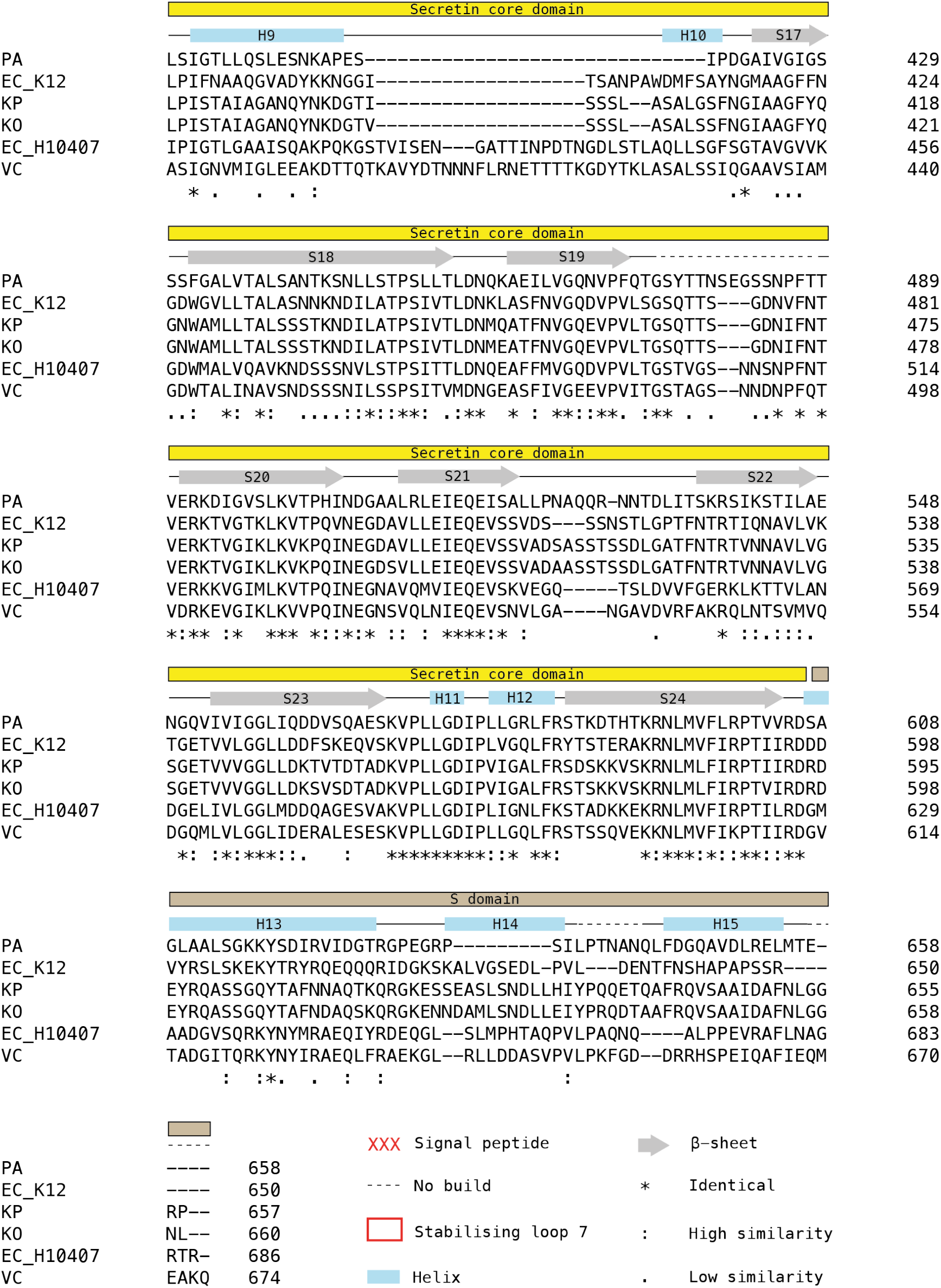
PulD secondary structure assignment and sequence alignment. Aligned PulD sequences include *Pseudomonas aeruginosa* (PA, Uniprot code P35818), *Escherichia coli* K12 (EC_K12, P45758), *Klebsiella pneumoniae* (KP, A0A0E1CJT4), *Klebsiella oxytoca* (KO, A0A0H3H6N4), *Escherichia coli* H10407 (EC_H10407, E3PJ86), *Vibrio cholerae* (VC, P45779).

**Figure S4.**
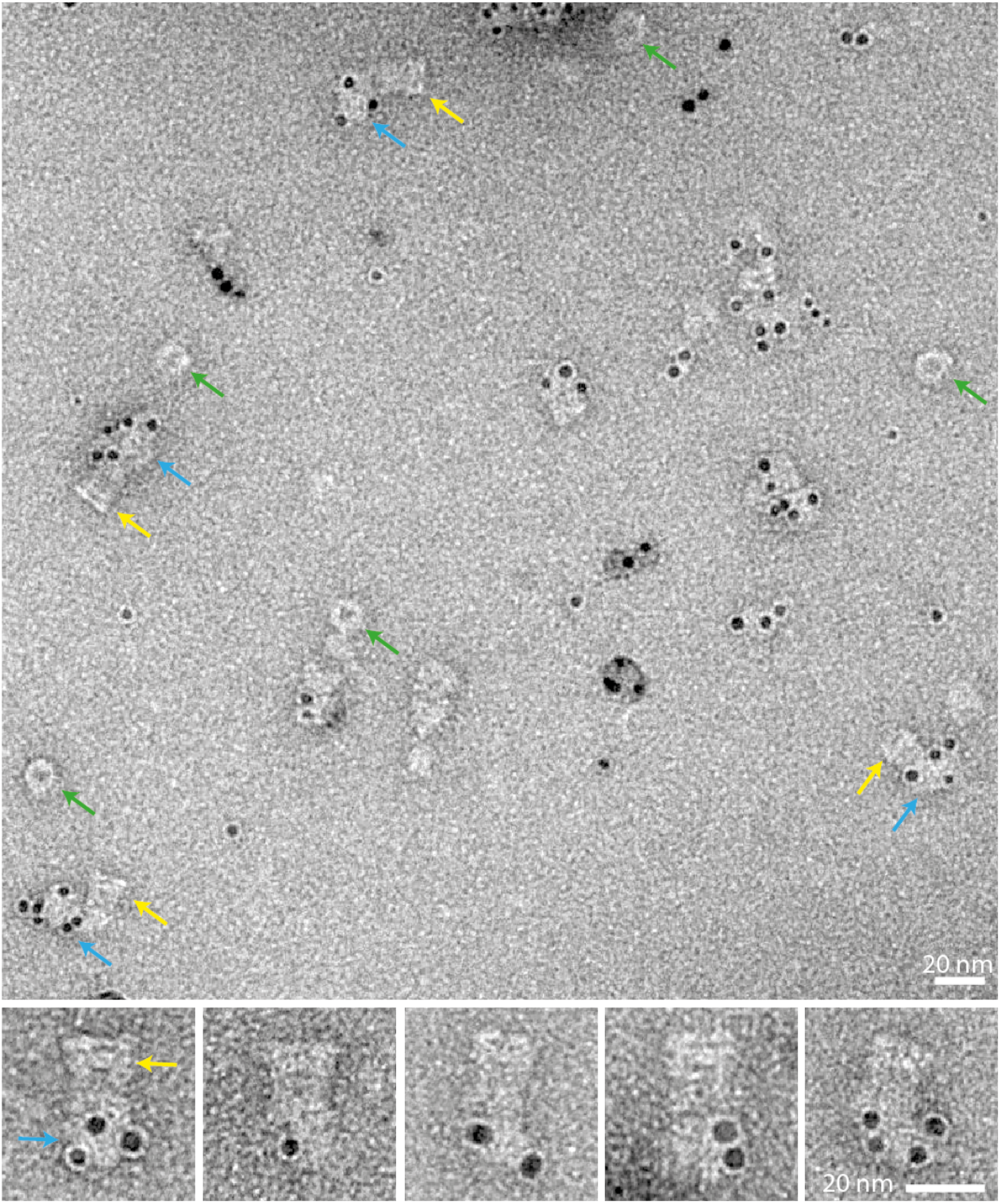
PulC N-terminus is located in the inner membrane assembly platform (AP). An overview negative stain EM image with a gallery of selected particles underneath. The PulC N-terminus was labelled with a hexahistidine tag at aa 61 within the Pul_CDELMNS_ complex. Ni-NTA gold beads localize to the AP (blue arrow). Gold beads did not localize to the secretin when attached to the AP (side view, yellow arrow) or when dissociated from the AP (top view, green arrow).

**Figure S5.**
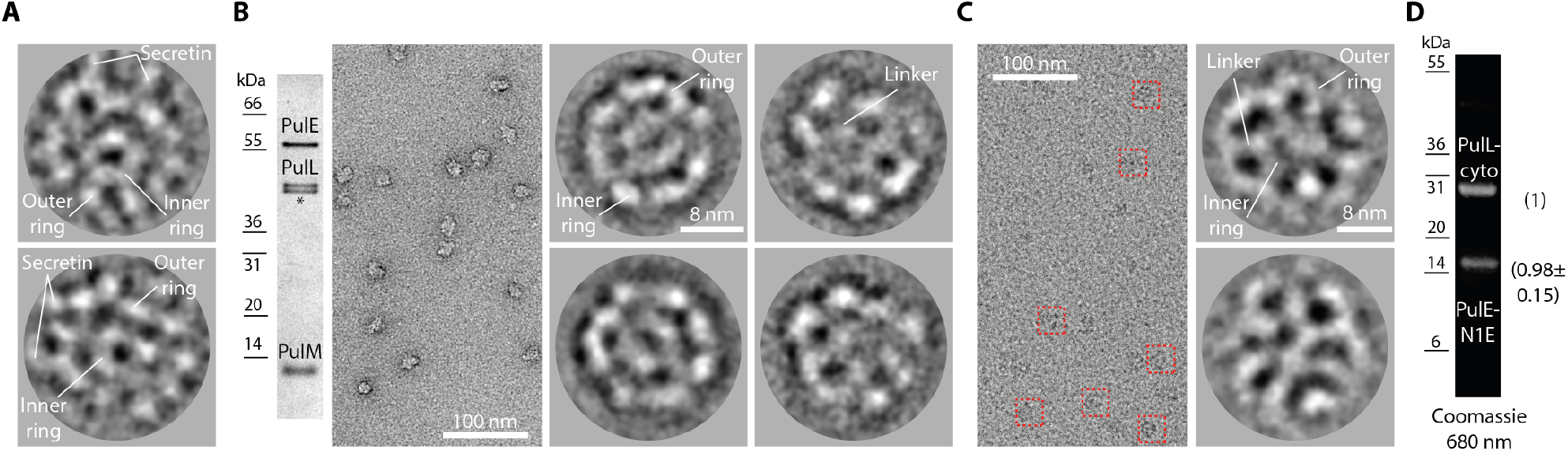
2D EM analysis of the Pul_CDELMNS_ assembly platform (AP) and the Pul_ELM_ complex. **(A)** Gallery of cryo-EM 2D class averages of the Pul_CDELMNS_ complex where the alignment and classification were focused on the AP only. A single preferred orientation was observed consistent with a bottom or end view. The base of the masked-out secretin is visible in the class averages. **(B)** GraFix purified Pul_ELM_ complex and EM analysis. (Left) Silver stain SDS-PAGE gel showing purified Pul_ELM_ complex. PulN bound weakly to the complex and was observed in only trace quantities after the GraFix ultracentrifugation step. (Right) Gallery of negative stain EM 2D class averages of the Pul_ELM_ complex. The same single preferred orientation was observed as in **A**. **(C)** Gallery of cryo-EM 2D class averages of the Pul_ELM_ complex. Note how the concentric ring ultrastructure is equivalent in both negative stain (as in **B**) and under cryo conditions. **(D)** SDS-PAGE of purified Pul_E-N1E/Lcyto_ complex. Fluorescent emission Coomassie R250 stained gel imaged at 680 nm with associated stoichiometry and standard deviation in parentheses. Stoichiometry measurements were determined from two independent purifications.

**Figure S6.**
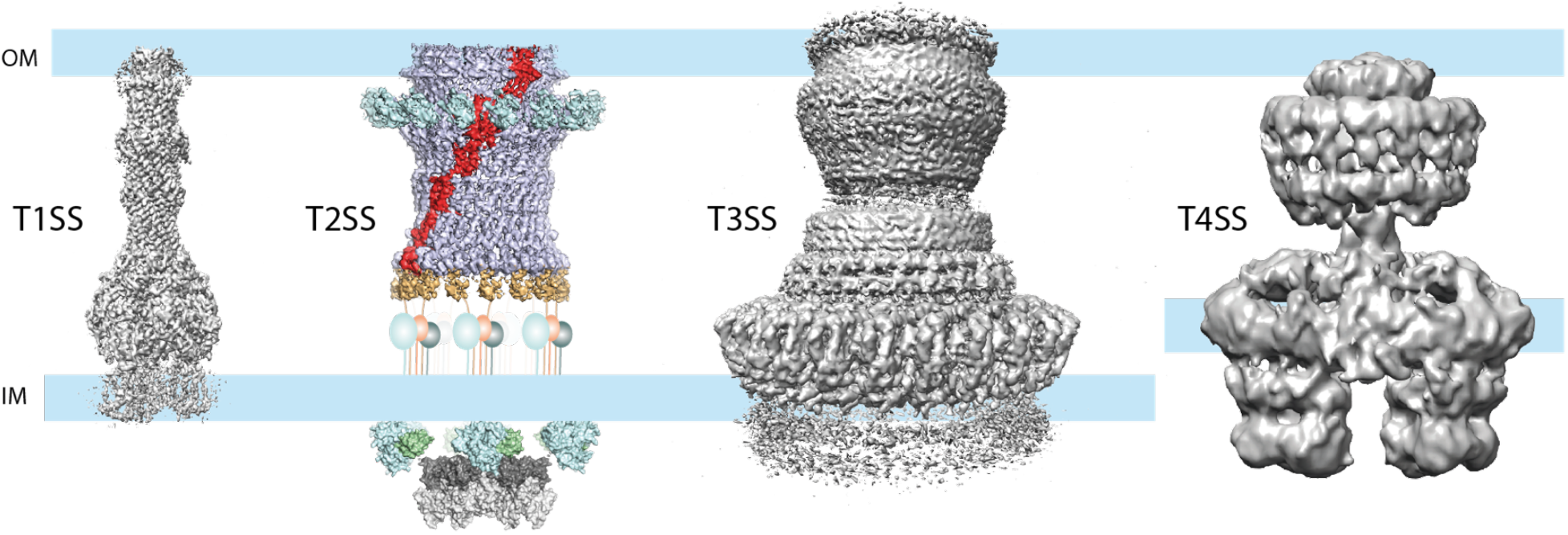
Comparison of bacterial type I-IV secretion systems. Substrate passes in an enclosed channel across the entire cell envelope in the type I (EMD 8636) and III (EMD 8400) secretion systems^38, 41^. In the T2SS, the substrate must enter the periplasm before outer membrane translocation through the secretin channel. In the type IV secretion system^36^, it is still unclear whether substrate passes across the cell envelope directly from the cytoplasm or is loaded first into the periplasm before being secreted.

**Supplementary Table 1.**
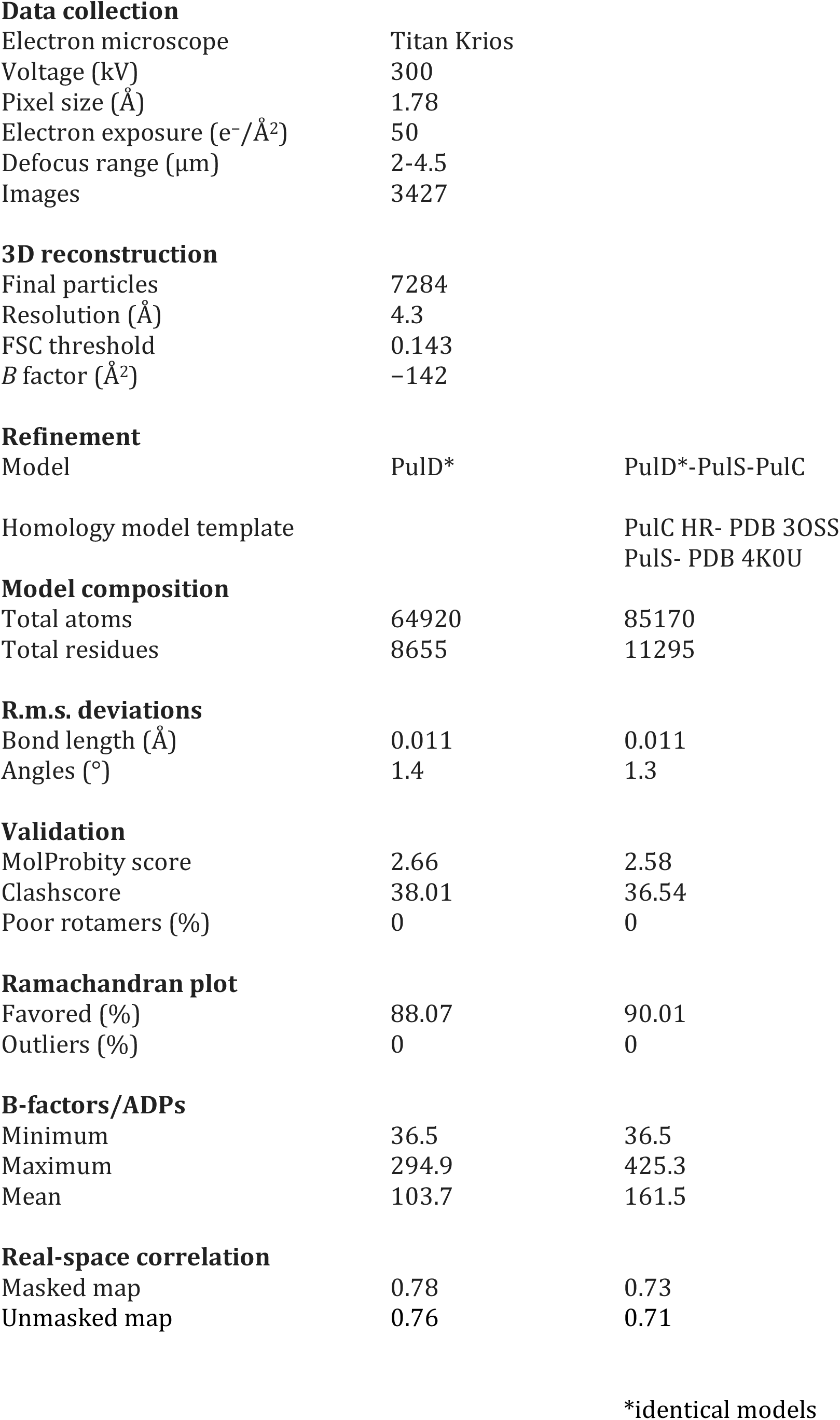
3D reconstruction and refinement statistics.

**Supplementary Table 2.**
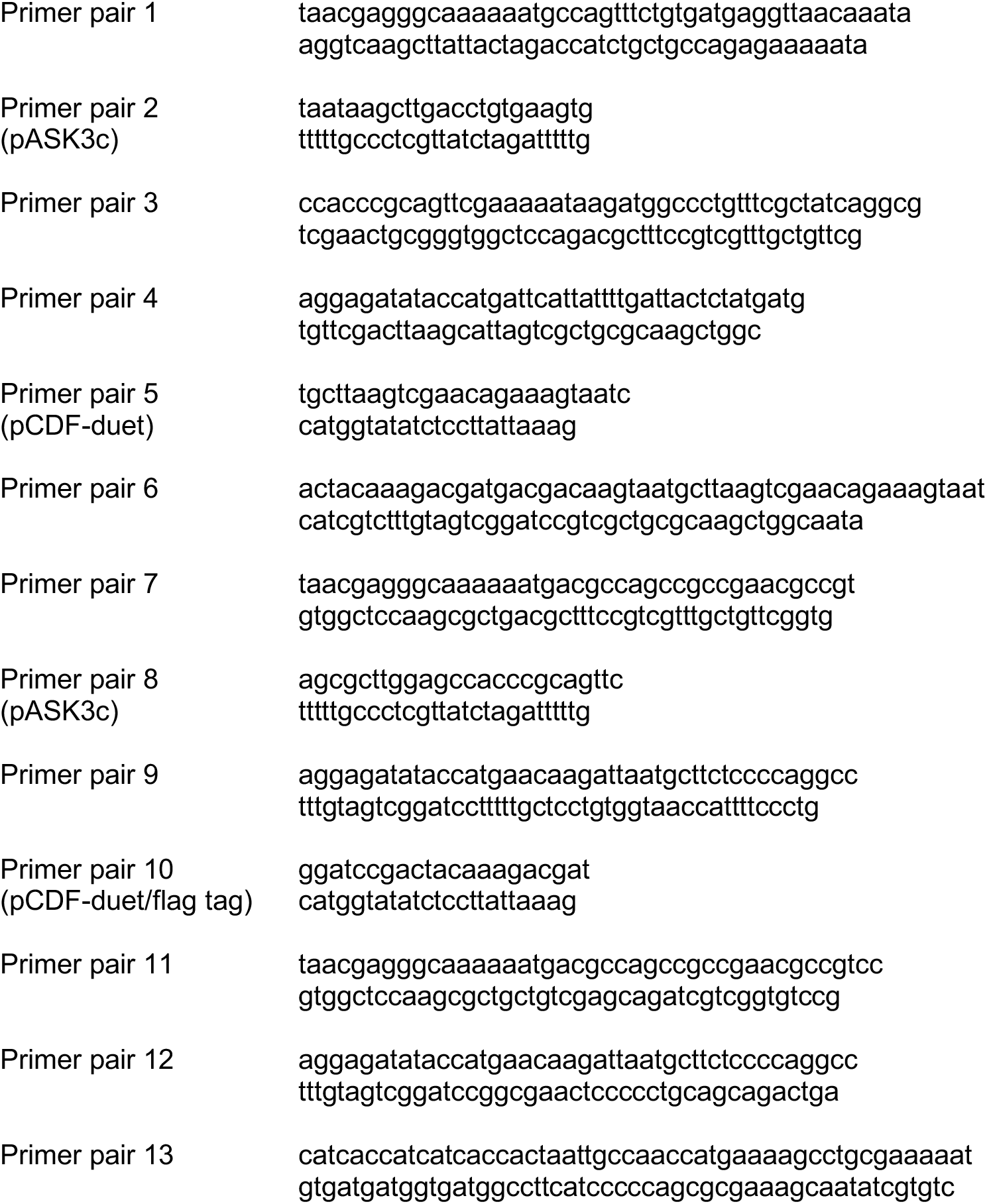
Cloning primers.

## Methods

### Cloning, protein expression and purification

All clones were generated using a modified version of the Gibson isothermal DNA assembly protocol^42^ where the *Taq* DNA ligase was omitted from the one-step isothermal DNA assembly. To obtain the Pul_CDELMNS_ complex, the *Klebsiella pneumoniae* T2SS operon encoding genes from *pulC* to *pulO* was cloned into pASK3c vector (IBA-GO) using primer pair 1 and 2 (for all primers used see Supplementary Table 2). A StrepII tag was then added at the C-terminus of PulE using primer pair 3. The *pulS* gene was cloned into pCDF-duet vector using primer pair 4 and 5. A C-terminal Flag tag was then inserted using primer pair 6. Uniprot codes for individual genes are as follows: *pulC* KPHS_08880, *pulD* KPHS_08870, *pulE* KPHS_08860, *pulF* KPHS_08850, *pulG* KPHS_08840, *pulH* KPHS_08830, *pulI* KPHS_08820, *pulJ* KPHS_08810, *pulK* KPHS_08800, *pulL* KPHS_08790, *pulM* KPHS_08780, *pulN* KPHS_08770, *pulO* KPHS_08760, *pulS* KPHS_08940. The clones were co-transformed into *Escherichia coli* C43 (DE3) electro-competent cells (Lucigen) modified here to incorporate a *pspA* gene knockout using a Lambda Red recombinase strategy^43^ (PspA is a common contaminant induced by PulD over-expression). Cells were grown on selective LB-agarose plates with chloramphenicol (30 µg/ml) and spectinomycin (50 µg/ml). 2xYT media was inoculated and cells grown at 37°C until induction at OD_600_ = 0.5-0.6 with anhydrotetracycline (AHT, 0.2 mg/L) and isopropyl β-D-1-thiogalactopyranoside (IPTG, 0.24 g/L). Cells were grown for ∼15 hr at 19 °C and processed immediately. Pellets were re-suspended in ice-cold buffer 50 mM Tris-HCl pH 7.5, 5 mM EDTA, treated with DNase I, lysozyme and sonicated on ice. The lysate was clarified by centrifugation at 16,000g for 20 min. The membrane fraction was collected by centrifugation at 142,000g for 45 min. Membranes were mechanically homogenized and solubilized in 50 mM Hepes-NaOH pH 7.5, 150 mM NaCl, 1 % w/v DDM (Anatrace) and 5 mM EDTA at room temperature for 30-40 min. The suspension was clarified by centrifugation at 132,000g for 15 min. The supernatant was loaded onto a StrepTrap HP column (GE Healthcare) and washed with 50 mM Hepes-NaOH pH 7.5, 150 mM NaCl, 0.06 % w/v DDM and 5 mM EDTA (Buffer W) at 4 °C. All prior buffers were supplemented with EDTA-free cOmplete protease inhibitor tablets (Roche). The protein sample was eluted in Buffer W supplemented with 2.5 mM desthiobiotin (IBA) but with protease inhibitors removed. Peak fractions were pooled, 0.05 % glutaraldehyde (Sigma-EM grade) added and incubated on ice for 10 min before quenching with 100 mM Tris-HCl pH 7.5. The sample was batch incubated with Flag resin (Sigma) for 1 hr. Flag resin was washed with Buffer W and then eluted with the same buffer supplemented with 3xFlag peptide. Peak fractions were collected and used immediately. LC-MS/MS confirmed the identity of bands identified by SDS-PAGE.

To obtain purified Pul_ELMN_ complex, the full-length *pulE* gene was cloned into pASK3c vector to include a C-terminal StrepII tag (primer pair 7 and 8). The *pulL*, *pulM* and *pulN* region of the T2SS operon was cloned into pCDF-duet vector with a PulN C-terminal Flag tag (primer pair 9 and 10). These vectors were co-transformed and the same initial purification strategy was then followed as for the Pul_CDELMNS_ complex with the exception that no glutaraldehyde or Tris quenching buffer were added subsequent to elution from the Strep column. Protease inhibitor tablets were included in all buffers. After elution from the Flag column, due to sample heterogeneity as judged by negative stain EM, GraFix^32^ was undertaken. Using Beckman Ultra-Clear 4.2 ml 11×60 mm ultracentrifugation tubes 2.1 ml of 50 mM Hepes-NaOH, 150 mM NaCl, 0.06 % w/v DDM, 30 % v/v glycerol, 5 mM EDTA and 0.1 % glutaraldehyde was loaded under 2.1 ml of the equivalent but with 10 % v/v glycerol. A continuous gradient was made using a BioComp Gradient Master cycle set for 66 seconds at 83° tilt and 22 rpm. An equivalent gradient was also made but omitting glutaraldehyde so that the sample could be analysed by SDS-PAGE after centrifugation. The Pul_ELMN_ sample was split in half and loaded onto the gradients ±glutaraldehyde. Samples were spun at 71,000g for 16 hr at 4°C using a Beckman Ti 60.1 swing rotor. 150 µl fractions were collected manually and analysed. Note that PulN was generally lost from the Pul_ELMN_ complex during GraFix yielding just Pul_ELM_.

To obtain purified Pul_E-N1E/Lcyto_ complex, the N1E domain comprising aa 1-108 from *pulE* were cloned into pASK3c vector to include a C-terminal StrepII tag (primer pair 8 and 11). For Pul_Lcyto_, aa 1-235 relating to the cytoplasmic domain of PulL were cloned into pCDF-duet vector with a C-terminal Flag tag (primer pair 10 and 12). The same 2-step affinity chromatography purification strategy was then followed as for PulELMN but excluding Grafix. Note that the PulE-N1E/Lcyto complex readily purifies from the membrane fraction despite the removal of the PulL trans-membrane domain.

### Gold labelling

This was performed on the Pul_CDELMNS_ complex modified to include a hexahistidine tag within PulC after aa 61 (primer pair 13). Purification was the same as for the Pul_CDELMNS_ complex but without the addition of fixative. Protease inhibitor tablets were included in all buffers. 5 nm Ni-NTA-Nanogold (Nanoprobes) pre-washed in Buffer W was added to 10 µl of the protein sample and incubated for 30 min at 4 °C. A homemade continuous carbon grid was deposited on the 10 µl sample for 3 min, blotted and washed 2 times in Buffer W supplemented with 10 mM imidazole, then 2 times in Buffer W before being stained with 3 drops of 2 % uranyl acetate.

### Stoichiometry determination

Pul_CDELMNS_ complex was purified as above with the exception that no glutaraldehyde or Tris quenching buffer were added subsequent to elution from the Strep column. Protease inhibitor tablets were included in all buffers. Six independent purifications were extracted using phenol to disrupt PulD multimerization^44^. Before extraction samples were divided and phenol treated in duplicate. The samples were precipitated with an equal volume of phenol and immediately vortexed. Four volumes of ice cold acetone were then added and vortexed again. The mixture was kept overnight at −20 °C. The precipitate was pelleted in a bench-top centrifuge at 14,000 g for 30 min at 4 °C, washed once with ice cold acetone, dried under vacuum and resuspended in loading buffer supplemented with 10 mM DTT. Both duplicates for each independent purification were analyzed by SDS-PAGE densitometry and quantification of fluorescent emission using both Coomassie Blue R250 and Sypro Ruby dyes. The Coomassie R250 procedure was as described^24^ but with extended stain and destaining times. Briefly, gels were rinsed with water and then stained with 0.01% Coomassie Blue R250 in 50% methanol and 10% acetic acid for 20 min. Gels were rinsed with 40% methanol and 7% acetic acid and then destained twice with the same solution for 20 min each time. Finally, gels were soaked in 2-3 times in water until fully destained, usually 1-2 hours. Gels were imaged on a Licor Odyssey Fc at 680 nm. Sypro Ruby gel staining was undertaken following the manufacturer’s instructions (Biorad). Gels were imaged on a Biorad Chemidoc XRS imaging system at 302 nm. Image Lab software (BioRad) and ImageJ were used to calculate band intensities.

To determine the stoichiometry of the Pul_E-N1E/Lcyto_ complex, two independent purifications were undertaken. Samples were run in duplicate on gels and then stained using the Coomassie Blue R250 procedure as outlined above. Gels were imaged on a Licor Odyssey Fc at 680 nm.

### Electron microscopy sample preparation and data collection

For outer membrane complex (OMC) structure determination, 4 µl of purified Pul_CDELMNS_ complex solution was incubated for 30 seconds on glow discharged homemade continuous thin carbon grids before vitrification in liquid ethane using a Vitrobot Mark IV (FEI). Data was collected at 300 kV on a Titan Krios (M02 beamline at eBIC Diamond, UK) equipped with a Gatan Quantum K2 Summit detector. Images were acquired at a magnification of 28,090 yielding 1.78 Å/pixel using EPU software. Images were dose-weighted over 40 frames with 12 second exposures. Total dose was ∼50 e/Å^2^.

All other cryo and negative stain datasets were collected in-house at 200 kV on a Tecnai F20 microscope equipped with Falcon II direct electron detector. For cryo data, Pul_CDELMNS_ complex (Figure 6B-C and Figure S5A) was vitrified as described above. The Pul_ELM_ GraFix treated sample required glycerol removal so that 4 µl of sample was loaded onto the glow discharged continuous carbon EM grid and after 1 min incubation was washed 4 times in Buffer W before plunge freezing. Images were acquired at a magnification of 90,909 yielding 1.65 Å/pixel using EPU software. Images were collected over 54 frames with 3 second exposures. Total dose was ∼50 e/Å^2^. For negative stain data, 4 µl of Pul_CDELMNS_ complex was loaded onto the glow discharged continuous carbon EM grid, after 40 seconds the grid was washed with 3 drops of distilled water and stained with 3 drops of 2 % uranyl acetate. Pul_ELM_ GraFix treated sample was similarly incubated on EM grids, washed iteratively with 4 drops of 15 µl Buffer W and then negatively stained as above. Images were acquired at a magnification of 90,909 using EPU software. Single frames were collected with 1 second exposure and total dose ∼15 e/Å^2^.

### Image processing

For OMC structure determination, individual movie image frames were aligned with MotionCor2 ^45^ and the contrast transfer function estimated using Gctf 1.06^46^. Low quality images were discarded and 3427 micrographs used for subsequent reconstruction in Relion 2.1^47^. Initial manual particle picking was focused on the OMC/secretin region of the Pul_CDELMNS_ complex. For particle extraction a box and mask diameter were chosen so that contributions from the inner membrane assembly platform (AP) were excluded. In this way, low-resolution 2D class averages of just the OMC were used as a template for auto-picking. OMC side views only were prevalent in this dataset, which provided a sufficiently even equatorial band distribution for a reliable reconstruction^48^. Low quality particles were removed by 4 rounds of 2D classification resulting in a stack of 36,240 particles. A single round of 3D classification was undertaken generating 10 classes. The *Vibrio cholerae* GspD reconstruction EMDB-1763 was used as an initial model filtered to 40 Å^49^. C15 symmetry was applied based on top views of the OMC (obtained in an alternative Pul_CDELMNS_ complex purification) and an unambiguous 15 peaks observed from the rotation auto-correlation function calculation (Figure 1C). A single class containing 7284 particles was used for the final refinement, which attained 4.4 Å resolution. Post-processing yielded 4.3 Å resolution with an auto-estimated B-factor ^50^ of −142 Å^2^ applied to sharpen the final 3D map for model building. A locally sharpened map was also generated using LocScale^39^ once initial models were built. Further particle polishing and 3D refinement did not yield a marked increase in resolution. Resolutions reported are based on gold standard Fourier shell correlations (FSC) = 0.143. Statistics for data collection and 3D refinement are included in Supplementary Table 1. Local 3D refinements with various particle subtraction strategies focusing on PulS or PulC HR domain did not markedly improve resolution.

To generate all other 2D class averages both in cryo and negative stain conditions as for Pul_CDELMNS_ and Pul_ELM_ complexes, the following protocol was followed. Working initially within Relion 2.0 or 2.1, Gctf 1.06 was used for estimating the CTF. Negative stain micrographs were phase flipped. Low quality micrographs were discarded. Initial 2D class averages were generated from a manually picked stack to yield templates for autopicking. 3-4 rounds of 2D classification were then undertaken to remove low quality particles. Using the ‘relion_stack_create’ a cleaned image stack was generated for further processing. For cryo-EM images the stack was created from phase flipped particles. In Imagic^51^, particles were normalised, band pass filtered, centred and subjected to reference-free MSA and classified. The best classes, typically judged by lowest variance, were used as references for multi-reference alignment (MRA) in Spider^52^ followed by MSA and classification in Imagic. This cycle of MRA and MSA was typically iterated a further 2-3 times. For negative stain Pul_ELM_ data, 2746 micrographs yielded 81,727 extracted particles and a cleaned stack of 63,014 particles. For cryo Pul_ELM_ data, 467 micrographs yielded 4937 hand-picked particles, and all particles were used for subsequent MSA and MRA. For Pul_CDELMNS_, 3059 micrographs yielded 89,381 extracted particles and a cleaned stack of 66,855 particles for subsequent MSA and MRA (Figure 6B). From this same dataset, 1148 micrographs were then used to hand-pick 4020 AP-focused particles (Figure 6C and Figure S5A). Rotation auto-correlation functions were calculated using Imagic.

### Model building

A PulD homology model was generated with I-Tasser^53^ using the *E. coli* GspD PDB 5WQ7 as a template. This yielded a starting model for the secretin core, and N3, N2 and N1 domains. *E. coli* GspD shares 57 % sequence identity with PulD. A homology model for the N0 domain was generated using Swissmodel^54^ with the relevant part of 3OSS as a template (51 % sequence identity). Homology models were rigid body fitted into the map using Chimera^55^ Fit in Map function. Using these models as a starting guide and the side chain detail from bulky residues to confirm sequence register, Coot^56^ was used to manually build a complete model for PulD aa 27-652 excluding aa 288-303, 462-470 and 632-637. The model was further refined using real-space refinement in Phenix^57^ with secondary structure, geometry and NCS restraints applied. For the low-resolution regions specific to the PulC HR domain, a homology model based on ETEC GspC HR domain PDB 3OSS (27 % sequence identity) was generated using Swissmodel. For fitting this homology model, the PDB 3OSS which includes the ETEC GspD N0 domain was first superimposed onto the PulD N0 domain. The PulC HR domain homology model was then superimposed onto the ETEC GspC HR domain from PDB 3OSS resulting in a near perfect fit within the map. The Chimera Fit in Map function was then applied to PulC HR domain resulting in a minor shift so that the PulC HR-PulD N0 domain complex has a RMSD C*α* = 1.4 Å when aligned to PDB 3OSS (Figure S2C). For the low-resolution regions specific to the PulD S-domain C-terminus (aa 638-652) in complex with the PulS pilotin, a homology model was generated using Swissmodel based on the equivalent structure from *Dickeya didantii* PDB 4K0U (>50 % sequence identity for both chains). The map was low pass filtered to 8 Å and the homology model initially fitted manually so that the PulS lipidated N-terminus orientated towards the membrane. The Chimera Fit in Map function was then used for final positioning. Cross-validations were carried out as previously described^10, 58^ using the auto-estimated B-factor sharpened map. Briefly, the PulD secretin model was displaced randomly by 0.2 Å and then refined against a map reconstructed from one of the independent data halves (Half map 1). FSC curves were then calculated using the resulting model and Half map 1 (FSC_work_). FSC curves were also calculated between this same model and another reconstruction generated from the other independent data half (Half Map 2 and FSC_free_). The similarity between FSC_work_ and FSC_free_ curves indicates an absence of overfitting within the PulD secretin model (Figure S1D). The final models were assessed using Molprobity^59^ and statistics outlined in Supplementary Table 1.

